# Comparative Analysis of Ultrafine Particulate Matter, Black Carbon, and Polystyrene Nanoplastics Identifies Mitochondrial Stress Adaptation as a Conserved Mechanism of Immunotoxicity

**DOI:** 10.64898/2026.07.29.741662

**Authors:** Pradyumna Kumar Mishra, Apoorva Chouksey, Aneha K Rajan, Vikas Gurjar, Ashwani Pathak, Aniket Aglawe, Ravi Prakash Tiwari, Debabrata Dash, Prakash Punj Dwivedi, Rajnarayan Tiwari, Devojit Kumar Sarma, Rupesh K. Srivasatava

## Abstract

Several studies have been conducted on human exposure to ultrafine particulate matter (UFPM), Black carbon (BC), and polystyrene nanoplastics (PS-NPs). However, it remains unclear whether different chemical types of environmental nanoparticles induce a similar mitochondrial stress response or a unique particle-specific response. In the present study, we examined the molecular mechanisms underlying nanoparticle-induced mitochondrial stress response and immunotoxicity using human peripheral blood mononuclear cells exposed to UFPM, BC, and PS-NPs under similar experimental conditions. Oxidative stress, mitochondrial adaptation, respiratory chain integrity, mitochondrial integrated stress response, inflammatory signaling, and systems-level interactions between molecules were analyzed through the evaluation of the expression of NRF2, HIF-1α, PGC-1α, TFAM, OMA1, DELE1, mitochondrial ND1, Complex I-V, NF-κB, TNF-α, and NLRP3 and the use of principal component analysis, hierarchical clustering, and correlation networks. All three nanoparticles caused oxidative stress and mitochondrial dysfunction with different kinetics and mechanisms. UFPM mostly induced an acute antioxidant response and mitochondrial adaptation; BC led to chronic mitochondrial dysfunction, chronic activation of the OMA1-DELE1-mediated mitochondrial ISR pathway, and inflammation; while PS-NPs induced low but chronic mitochondrial adaptation along with mitochondrial biogenesis and stress responses. Our systems-level analysis showed that oxidative stress, mitochondrial adaptation, mitochondrial ISR, and inflammation represent a highly connected molecular network regardless of the physicochemical nature of the nanoparticles, with the OMA1- DELE1 axis being a key regulatory node connecting mitochondrial stress response and inflammation. Overall, we have found that mitochondrial stress response is a common mechanism underlying the toxicity of chemically different nanoparticles and have also revealed particle-specific stress-response dynamics responsible for the degree and persistence of cellular damage. The current work presents novel insights into the molecular mechanisms of nanoparticle-induced immunotoxicity and suggests OMA1, DELE1, NRF2, PGC-1α, TFAM, ND1, and Complex I-V as potential biomarkers.

## 1. Introduction

Humans are continuously exposed to nanoscale-sized particles, including ultrafine particulate matter (UFPM), Black carbon (BC), and polystyrene nanoplastics (PS-NPs). Such particles are found abundantly in current levels of pollution and have been identified as potential sources of immunotoxicity and inflammation. Even though they exist together in the real-world exposure settings of urban air, indoor spaces, and occupational sites, they are chemically unique molecular exposure paradigms, which differ substantially in environmental source, chemical composition, surface chemistry, physicochemical properties, cellular uptake and intracellular transport, as well as biological durability (Calderón-Garcidueñas & Ayala, 2022; Wang et al., 2024). With increasing knowledge of their harmful impacts on human health, most studies have explored UFPM, BC, or PS-NPs separately, employing diverse biological models and endpoints. It is unclear whether the unique chemistry of these nanoparticulate materials influences the universality of the mitochondrial stress response pathway or whether they elicit specific adaptive responses that result in differing biological effects despite similar exposures.

Through redox cycling, UFPM causes toxicity and Fenton reactions via transition metal ions, which drive oxidative stress, high ROS production, and mitochondrial damage (Li et al., 2003; Bhargava et al., 2017). BC, on the other hand, consists of graphitic carbon with PAHs and transition metals whose high chemical stability contributes to cell uptake and subsequent cellular events such as lysosomal dysfunction, metabolic changes, and chronic inflammation (Schraufnagel et al., 2019). The toxicity of NPs is associated with the rapid formation of a dynamic biomolecular corona responsible for receptor-mediated entry and intracellular transport, organelle targeting, and activation of immune pathways (Gopinath et al., 2021). Despite the various upstream events that initiate cellular stress induced by the three types of environmental nanoparticles, diverse physicochemical signals are channeled into mitochondria, the organelle responsible not only for cell bioenergetics but also for integrating metabolic, redox, and inflammatory signaling.

Though there is growing awareness about the importance of mitochondrial dysfunction in the development of nanoparticle toxicity, the existing knowledge base has been established using an isolated framework of research for individual nanoparticles. Whereas UFPMs have been widely studied for oxidative stress, mitochondrial DNA damage, and mitochondrial epigenetics (Byun et al., 2013), black carbon has been widely studied for mitochondrial dynamics, mitophagy, and quality control (Shang et al., 2022). On the other hand, research on NPs has mainly revolved around mitochondrial unfolded protein response signaling and adaptive mitochondrial remodeling (Lee et al., 2022). Though these studies have contributed immensely to expanding the knowledge base about individual nanoparticle types, they have focused only on specific aspects of mitochondrial biology under varying experimental conditions. These results lead to an important research gap concerning whether chemically unique nanoparticulate environmental contaminants utilize the same mitochondrial pathway or follow distinct stress-induced pathways that result in bioenergetic dysfunction and immunotoxicity. Answering such questions requires a system-wide investigation that can concurrently explore oxidative stress, mitochondrial adaptations, mitochondrial genomic integrity, mitochondrial respiration, and inflammatory signaling. To the best of our knowledge, there is no direct comparison of UFPM, BC, and NPs under similar exposure conditions using a comprehensive set of markers for mitochondrial stress, bioenergy, inflammation, and systemic responses.

The adaptive response of mitochondria to nanoparticles in the environment exists on a continuum from adaptation through bioenergetic dysfunction to inflammation induction, depending on the extent of mitochondrial damage. Mitochondrial dysfunction starts with increased ROS generation, redox imbalance, calcium accumulation, loss of mitochondrial membrane potential (Δψm), and impaired oxidative phosphorylation. These events collectively constitute an acute attack on mitochondria, initiating stress response mechanisms rather than causing immediate damage (Bhargava et al., 2019; Sharma et al., 2020; Cameron et al., 2022). First, acute mitochondrial injury initiates processes that help mitochondria repair themselves. One of the major elements of these processes is modulation of the HIF-1α-PGC-1α-TFAM pathway.

Deficiency of these adaptation mechanisms leads to an irreversible switch between reversible mitochondrial stress and persistent malfunction. When these mechanisms fail, Δψm is disrupted, oxidative phosphorylation malfunctions, and oxidative mtDNA damage occurs. In this situation, retrograde signaling and OMA1-DELE1-mediated ISR take place through which the organism adapts to mitochondrial stress (Guo et al., 2020; Fessler et al., 2020; Mishra et al., 2022). However, despite the recent identification of the OMA1-DELE1-ISR axis as an important coordinator of mitochondrial stress adaptation, whether distinct nanoparticles differentially engage the pathways has not been determined. Inflammation caused by the activation of NF-kB and the creation of the NLRP3 inflammasome is caused by mitochondria-induced stress; when the latter is unable to do this, it serves as the connection between mitochondrial injury and immunotoxicity (Grazioli & Pugin, 2018; Lyu et al., 2023; Mun et al., 2025; Zhang et al., 2025). Hence, mitochondrial stress can be considered a continuum ranging from adaptation to inflammation, suggesting that UFPMs, BC, and NPs induce mitochondrial stress through different pathways that converge on the same organelle.

However, if the extent of oxidative stress exceeds mitochondrial adaptability, dysfunctional processes extend to all components of the respiratory chain, resulting in defective oxidative phosphorylation, ATP production, and mitochondrial bioenergetics. Complexes I-V work in coordination to support electron transfer and ATP production, but disruption of these complexes leads to electron leakage, production of large amounts of ROS by the mitochondria, mitochondrial depolarization, and activation of mitochondrial stress pathways (Murphy, 2009; Okoye et al., 2023). Since the genes encoding respiratory chain proteins and complexes are present in the mitochondrial genome, particularly complex I, oxidative damage results in mitochondrial genotoxicity and aggravation of respiratory dysfunction (Ide et al., 2001; Scarpulla, 2011). Hence, determining respiratory chain function, along with assessing transcriptional stress pathways, provides a more thorough evaluation of nanoparticle-induced mitochondrial dysfunction.

Peripheral blood mononuclear cells (PBMCs) serve as a biologically relevant ex vivo system for understanding the mechanisms by virtue of the fact that they represent the initial circulating immune cells that come into contact with translocated environmental nanoparticles and depend greatly on the mitochondria-mediated metabolism of activation, cytokine production, and inflammatory response (Dobrovolskaia & Afonin, 2020; Soni et al., 2025). Thus, we established an exposure comparison model in which PBMCs were subjected to different environmentally relevant nanoparticles with unique chemical compositions under similar experimental conditions to study whether common or particle-specific mitochondrial stress adaptation pathways exist. We conducted a systematic analysis of mitochondrial adaptive signaling pathways (HIF-1α-PGC-1α-TFAM), mitochondrial stress signaling (OMA1-DELE1-ISR), respiratory chain integrity (Complex I-V and mtDNA), inflammatory signaling (NF-κB/NLRP3), and systems biology markers of these biological processes in PBMCs exposed to chemically distinct environmental nanoparticulate materials. We hypothesized that although UFPM, BC, and NPs affect mitochondrial homeostasis, each nanoparticle triggers unique mitochondrial stress adaptation pathways that lead to distinct molecular profiles and immunotoxicity.

## 2. Materials and methods

PBMCs were treated with UFPM, BC, and PS-NPs (1 µg/mL) for 6, 12, and 24 hours, respectively, following Institutional Ethics Committee clearance (Salinas et al., 2020; Adamiak et al., 2025). Oxidative stress and Δψm in mitochondria were assessed using flow cytometry, whereas oxidative DNA damage was also confirmed using the Fpg assay (Soni et al., 2025). The mRNA levels of oxidative stress-responsive genes, mitochondrial biogenesis, and inflammatory pathways (NRF2, SOD2, HIF-1α, PGC-1α, TFAM, RELA, NFKB1, and NLRP3) were analyzed by one-step quantitative real-time PCR; however, OXPHOS genes were quantified by probe-based quantitative real-time PCR. Mitochondrial DNA content and ETC complex activity were assessed using quantitative real-time PCR and activity measurements, respectively (Mishra et al., 2014; Bhargava et al., 2019). Immunotoxicity was determined by measuring IL-1β, IL-6, IL-8, and TNF-α levels by ELISA (Bhargava et al., 2017). Principal Component Analysis, K-means Clustering, and Pearson Correlation Coefficient were done using Python (Kaur et al., 2025). The experimental methods are detailed in the supplementary materials (Supplementary methods).

## 3. Results

### 3.1. Particle-specific temporal regulation of mitochondrial stress, bioenergetic, and inflammatory pathways

To evaluate the differential molecular responses generated by environmentally ubiquitous nanoparticles with chemically diverse properties, the temporal expression of genes implicated in mitochondrial stress signaling, oxidative stress, mitochondrial bioenergetics, and inflammatory signaling in PBMCs exposed to UFPM, BC, and PS-NPs for 6, 12, and 24 h (Figure 1) was investigated. Exposure of PBMCs to the three nanoparticles resulted in differential transcriptional remodeling among nanoparticles. In relation to the mitochondrial ISR, the transcriptional activation of genes such as OMA1 and DELE1 was consistently upregulated upon BC exposure, whereas UFPM and PS-NP exposures induced progressive and transient induction of gene expression, respectively. In addition, the antioxidant genes NRF2 and SOD2 were persistently activated in PBMCs exposed to BC, but UFPM and PS-NP exposures resulted in delayed and reduced antioxidant responses, respectively. Mitochondrial adaptation and bioenergetic genes were also differentially regulated depending on the nanoparticles used. While UFPM strongly induced HIF-1α at 12-24 h, it was not upregulated in cells treated with PS-NPs. In addition, PGC-1α was recovered from the suppression induced by UFPM, but PGC-1α was relatively constant in cells exposed to BC. BC and UFPM moderately upregulated TFAM, but TFAM was induced maximally in cells treated with PS-NPs after 24 h exposure. The mitochondrial respiratory gene ND1, on the other hand, remained stable when exposed to BC, while it was progressively inhibited when exposed to PS-NPs but restored after exposure to UFPM. Likewise, temporal variations in the inflammatory response were noted, with BC consistently upregulating the inflammation-related mediators TNF-α and NLRP3. On the other hand, UFPM showed high and transient expression of NF-κB, NF-κB p65, and TNF-α, where maximum induction occurred after 12 hours, followed by continuous induction of NLRP3. However, PS-NPs induced the weakest inflammatory response, characterized by low TNF-α expression and suppression of NF-κB signaling and NLRP3 activation. To determine whether the transcriptional alterations observed were functional consequences for the mitochondria, mitochondrial respiratory chain activities (Complexes I-V) were investigated after nanoparticle exposure (Supplementary Figs. S1 and S2). The changes in respiratory chain complex activities were nanoparticle exposure-dependent and supported the changes in transcription and mitochondrial oxidative phosphorylation.

**Figure 1.**
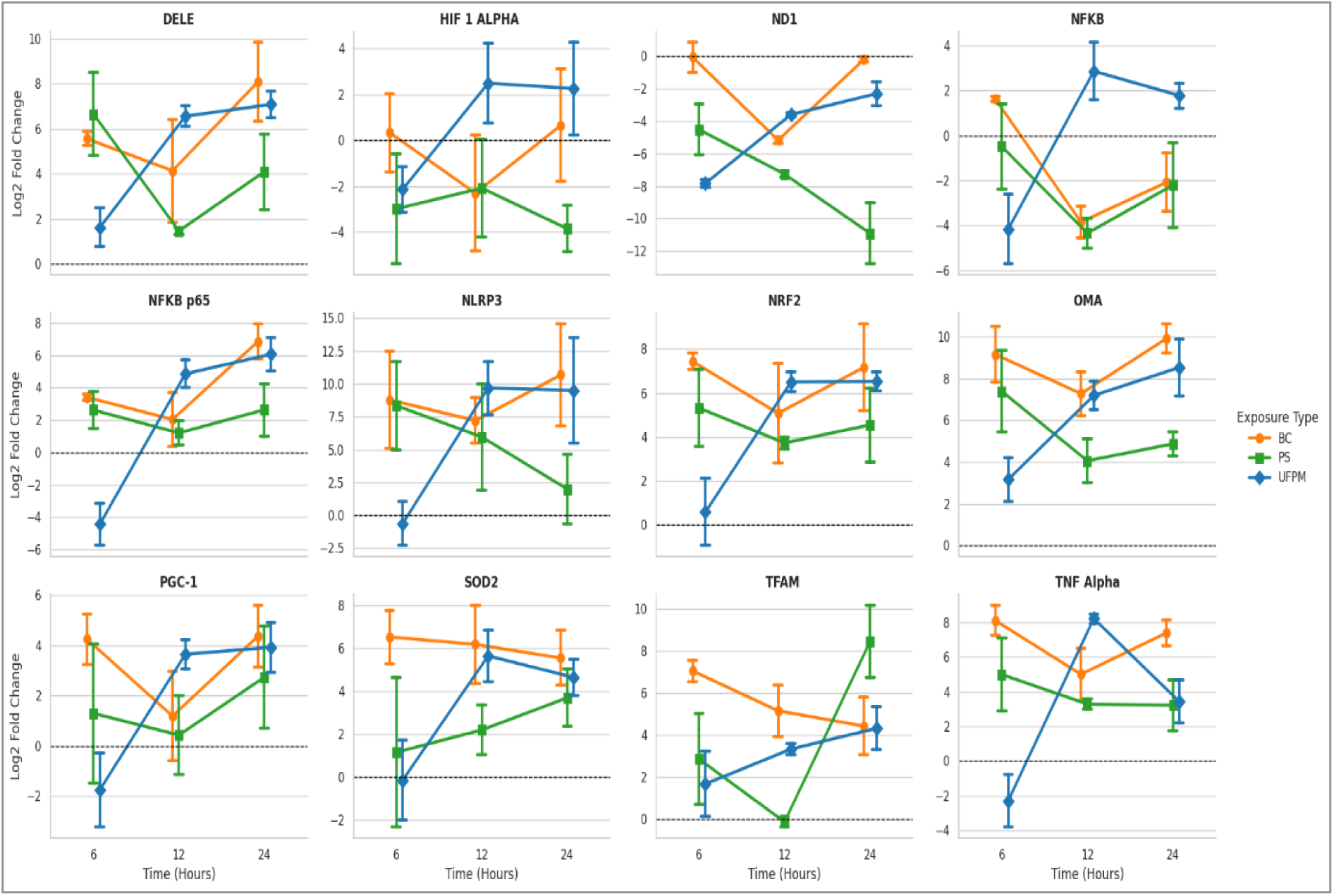
Time-dependent regulation of mitochondrial, inflammatory, and oxidative stress response gene expression following treatment with BC, PS-NPs, and UFPM. Time-dependent regulation of the expression of mRNAs for DELE, HIF-1α, ND1, NF-κB, NF-κB p65, NLRP3, NRF2, OMA1, PGC-1α, SOD2, TFAM, and TNF-α was examined at 6, 12, and 24 h post-treatment with black carbon (BC; orange), polystyrene nanoplastics (PS; green), and ultrafine particulate matter (UFPM; blue). The expression was measured relative to the respective internal control gene and presented as a log2 ratio compared to the untreated control sample, indicated by the horizontal dotted line at zero. Values are presented as mean ± SEM. The error bars show the standard error of the mean. The current time course experiment highlights time-dependent regulation of pathways related to mitochondrial stress response (DELE1-OMA1), hypoxia (HIF-1α), mitochondrial function (ND1, TFAM), oxidative stress (NRF2, SOD2), mitochondrial biogenesis (PGC-1α), and inflammation (NF-κB, NF-κB p65, NLRP3, and TNF-α).

Overall, the comparisons among the three particulates demonstrated the similarities and differences in molecular responses. While having distinct physicochemical properties, UFPM, BC, and PS-NPs converge on a common mitochondrial stress pathway that includes activation of the mitochondrial ISR, antioxidant defense, mitochondrial bioenergetic adaptation, and inflammatory signaling. Nevertheless, the magnitude and kinetics of activation were different for various particles. UFPM caused the most comprehensive remodeling of transcription over time, including rapid activation of adaptive and inflammatory responses. On the other hand, BC resulted in the most lasting activation of ISR and inflammatory responses, along with oxidative stress in mitochondria. In addition, PS-NPs induced the weakest inflammatory responses but the most prominent regulation of mitochondrial adaptive and biogenesis-related genes, including TFAM, as well as persistent suppression of ND1. Therefore, these results suggest that differentially physicochemical nanoparticles activate the conserved mitochondrial stress network together with the particle-specific pathway regulation, which determines differences in bioenergetic adaptation and inflammation.

### 3.2. Principal component analysis reveals exposure- and time-dependent transcriptional signatures

To assess the transcriptional relationship between the tested conditions (UFPM, BC, PS-NPs), PCA was applied to PBMCs exposed to UFPM, BC, and PS-NPs for 6, 12, and 24 h (Figure 2). Collectively, PC1 and PC2 accounted for 73.1% of the variation, with PC1 explaining 60.9% and PC2 12.2%, indicating that most of the variation can be explained by these two components. Clustering based on nanoparticle type demonstrated clear separation of the samples into different clusters, confirming the specificity of the molecular responses of PBMCs to each nanoparticle and the existence of common molecular responses. The progression of transcriptional alterations in the samples exposed to UFPM, BC, and PS-NPs for 6, 12, and 24 h was also seen. In general, samples harvested 24 h apart were located at separate positions compared to the samples obtained earlier. Moreover, of all the nanoparticle types, UFPM revealed the largest area covered by PCA projection with respect to time of exposure. BCs formed the smallest cluster with gradual movement in time. Samples obtained from PS-NPs formed more dispersed clusters in the PCA space compared to others, indicating greater variance in the response to PS-NPs over time.

**Figure 2.**
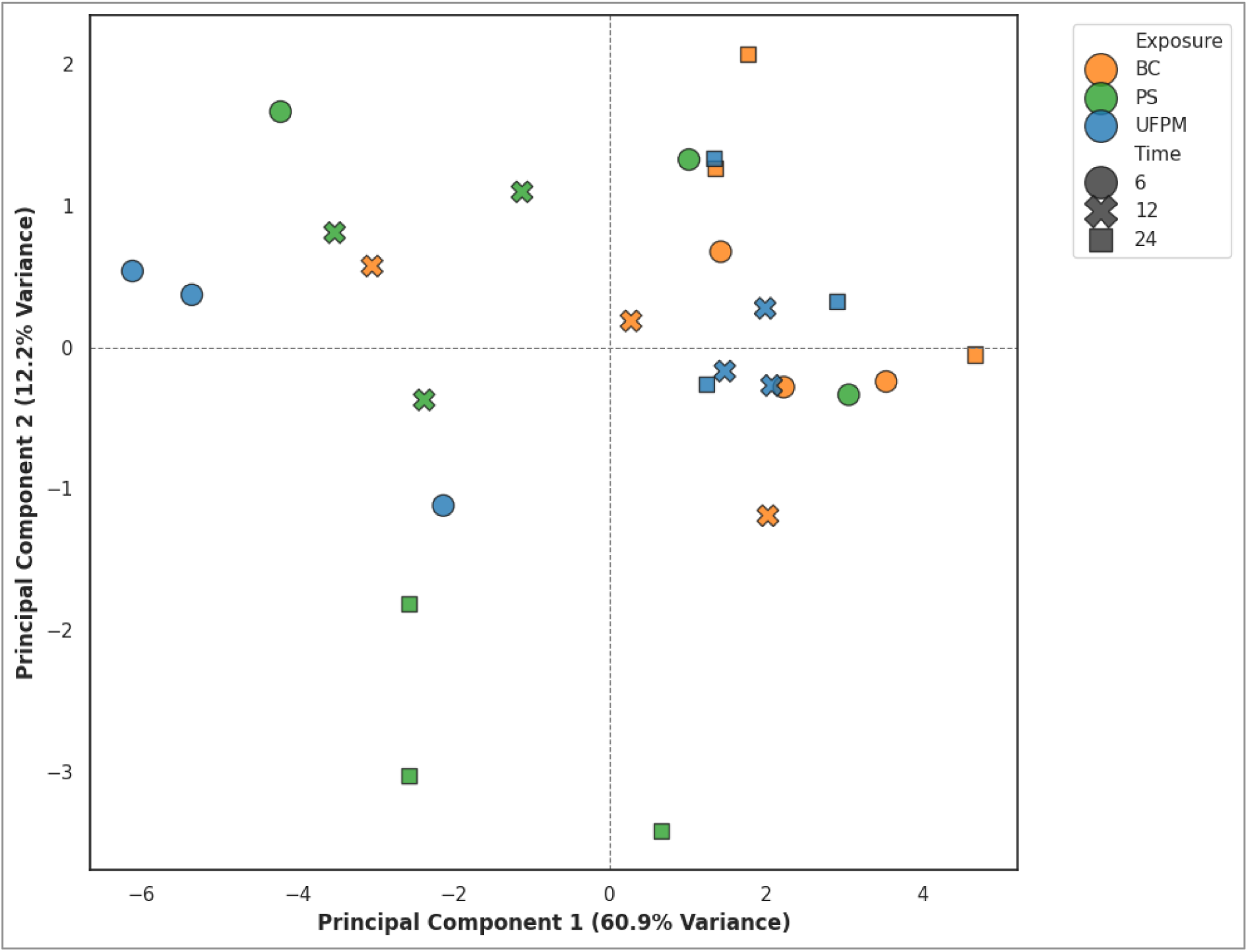
Principal component analysis (PCA) of transcriptional response upon exposure to BC, PS-NPs, and UFPM. A PCA was conducted using the normalized expression profiles of DELE1, HIF-1α, ND1, NF-κB, NF-κB p65, NLRP3, NRF2, OMA1, PGC-1α, SOD2, TFAM, and TNF-α following exposure to BC, PS-NPs, and UFPM for 6, 12, and 24 h. Color codes differentiate between types of exposures (BC - orange; PS - green; UFPM - blue), while different symbols differentiate between the time of exposure (circles - 6 h; crosses - 12 h; squares - 24 h). PC1 and PC2 account for 60.9% and 12.2% of total variance, respectively.

### 3.3. Differential temporal regulation of FAPY following particulate exposure

To determine the kinetics of mitochondrial oxidative DNA damage and the DNA repair response, the expression pattern of FAPY was compared between particulate-exposed and untreated PBMCs following BC, PS-NP, and UFPM exposure (Figure 3). Each nanoparticle type elicited unique kinetics of FAPY expression due to differences in exposure effects. While BC induced a gradual rise in FAPY expression in exposed cells, with mild up-regulation at 6 and 12 hours and the highest expression at 24 h, the untreated control showed the opposite trend, with down-regulation throughout the experiment. In contrast, FAPY was generated by PS-NP at 6 h and declined further at 12 and 24 h; however, it remained significantly higher than basal levels even at 24 h in the presence of the treated cells. UFPM exhibited another distinct time-dependent trend. FAPY expression was first downregulated at 6 h, nearly at baseline at 12 h, and reached the peak value only after 24 h of exposure, providing evidence of delayed activation of the DNA repair response. In contrast, the untreated UFPM samples demonstrated continuous down-regulation. Thus, the comparison of the expression of FAPY for the three particulates demonstrated distinctive kinetics of regulation depending on their nature. Although the three nanoparticles activate the DNA repair response, the timing and intensity of the response are determined by the physicochemical properties of exposure.

**Figure 3.**
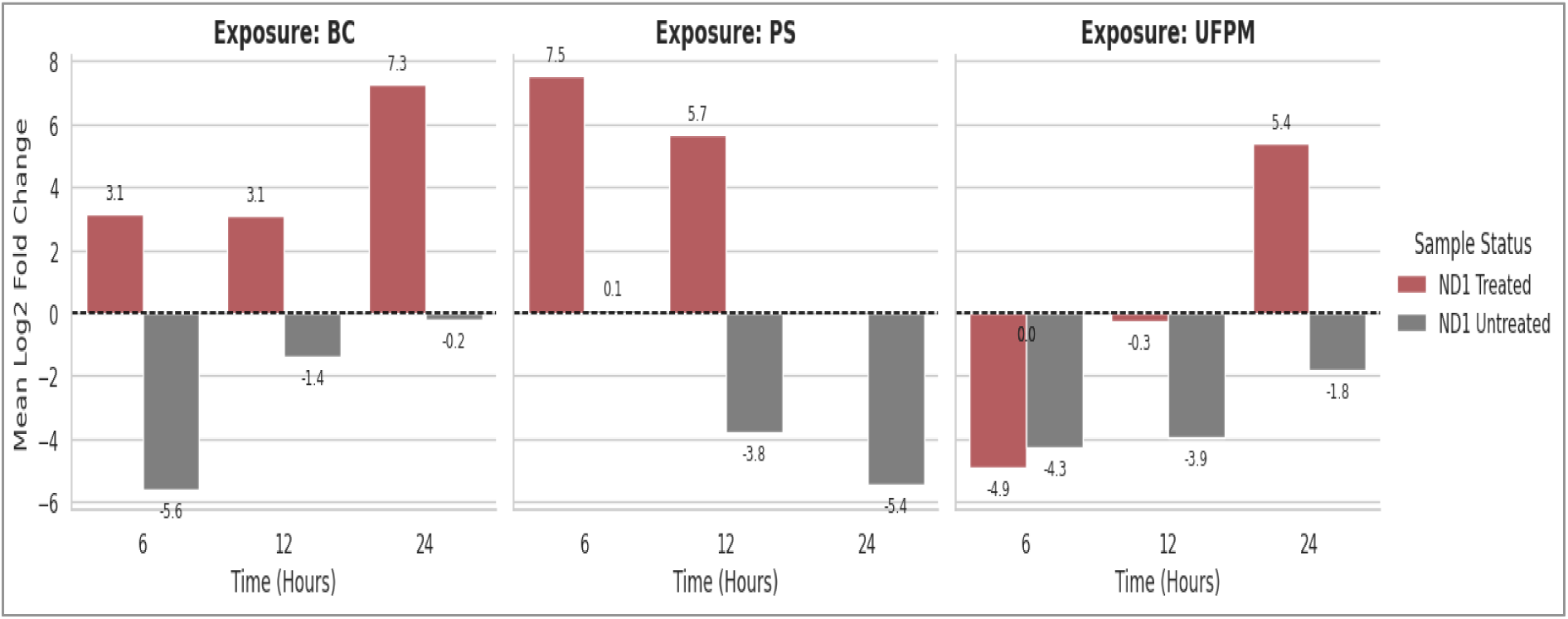
Time course of mitochondrial ND1 expression in treated and untreated cells exposed to BC, PS-NPs, and UFPM. Comparison of mean log2 fold change in mitochondrial ND1 expression between treated and untreated cells upon exposure to BC, PS-NPs, and UFPM for 6, 12, and 24 h was performed. The data shown represent the mean gene expression value obtained from Experiments 1 and 2. Values above zero indicate up-regulation, while those below indicate down-regulation relative to the untreated control cells marked with a horizontal dashed line (0). Bars are grouped according to exposure type and sampling time to facilitate comparison of treatment-dependent temporal changes in mitochondrial gene expression.

### 3.4. Two-way ANOVA identifies exposure- and interaction-dependent regulation of mitochondrial stress pathways

Two-way ANOVA was conducted to assess the independent and interaction effects of nanoparticle exposure and exposure time on gene expression (Figure 4). Of the three factors considered, the interaction effect of exposure × time appeared to have the strongest influence on the transcriptional profile, with significant interactions detected for ND1, NF-κB, NF-κB p65, TNF-α, NRF2, TFAM, DELE1, and OMA1 (all p < 0.05). The obtained data suggest that the transcriptional regulation of mitochondrial bioenergetics, oxidative stress, mitochondrial ISR, and inflammatory signaling depends on the combined effect of nanoparticle type and exposure time, rather than on each individual factor. Separate main-effects analyses showed that exposure affected the transcriptional activity of the ND1, TNF-α, SOD2, and OMA1 genes, while time had a significant effect only on the NF-κB p65 and TFAM genes. At the same time, the genes of NLRP3, HIF-1α, and PGC-1α did not have a significant main effect of either exposure or time, which means that their transcriptional regulation is mostly associated with coordinated pathways’ responses. Thus, mitochondrial respiration (ND1), mitochondrial ISR (OMA1-DELE1), antioxidant defense (NRF2-SOD2), mitochondrial biogenesis (TFAM), and inflammatory signaling (NF-κB/NF-κB p65 and TNF-α) are controlled by distinct exposure, time, and their interaction.

**Figure 4.**
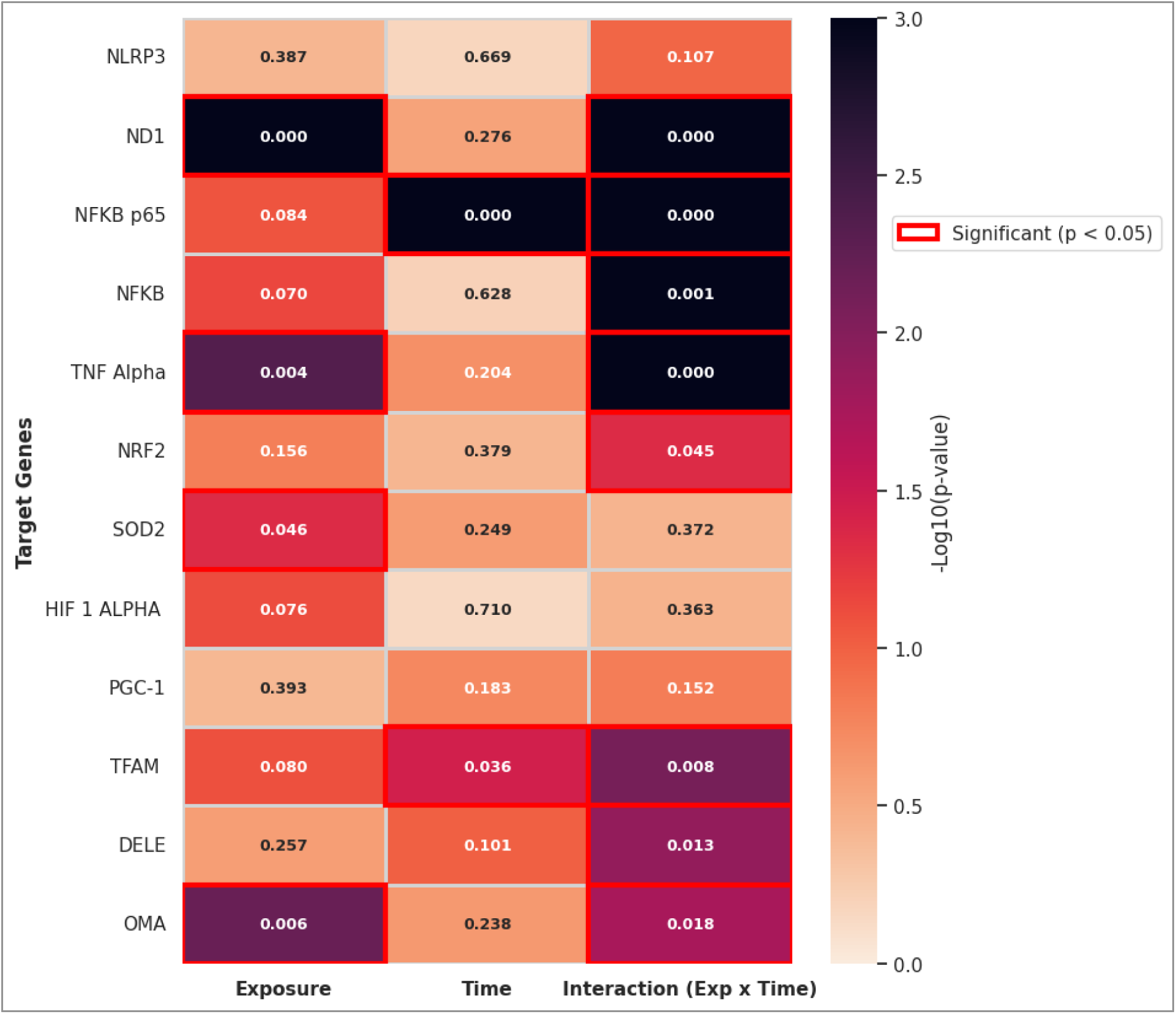
Two-way ANOVA reveals the impact of the interaction between exposure and time on the expression of mitochondrial stress-related genes, inflammatory genes, and oxidative stress-related genes. The heatmap represents the statistical significance of exposure, time, and the interaction between exposure and time on the expression of genes of interest using a two-way ANOVA test. Color in the cells denotes −log₁₀(p-value), where darker colors signify higher statistical significance. The values written in each cell are the corresponding p-values. Statistically significant impacts are red-framed (p<0.05). The test was conducted for NLRP3, ND1, NF-κB p65, NF-κB, TNF-α, NRF2, SOD2, HIF-1α, PGC-1α, TFAM, DELE1, and OMA1.

### 3.5. Hierarchical clustering reveals coordinated regulation of mitochondrial stress pathways

The hierarchical cluster analysis was conducted to understand the interaction of the different exposure groups with genes associated with mitochondrial stress, bioenergetics, oxidative stress, and inflammation (Figure 5). Heat map analysis showed different clusters of samples by both nanoparticle types and time of exposure, indicating progressive remodeling of transcriptional patterns with increasing duration of exposure. Generally, samples collected 24 h after exposure were separate from the corresponding 6 h and 12 h samples and had higher expression of stress-induced genes. In contrast, early exposure groups had relatively low gene expression, whereas the 12 h samples had intermediate transcriptional profiles, corresponding to the transition from the initial adaptive response of mitochondrial stress. The gene dendrogram analysis also showed two major modules of co-expression. First, the module was composed of ND1, HIF-1α, and NF-κB genes that encode proteins involved in mitochondrial respiratory function, hypoxic adaptation, and inflammatory signaling, respectively. The second module included NLRP3, OMA1, NF-κB p65, PGC-1α, TFAM, SOD2, TNF-α, DELE1, and NRF2, which indicates the involvement of mitochondrial ISR, oxidative stress, antioxidant defense, mitochondrial biogenesis, and inflammatory signaling. Prolonged exposure to BC and UFPM was defined by elevated activity of genes involved in mitochondrial stress and inflammation, such as OMA1, DELE1, NRF2, TNF-α, and NLRP3, in contrast to the early exposure groups, in which a decrease in transcriptional activity was observed. In summary, the results of hierarchical clustering indicate that the transcriptional response to nanoparticles is organized into coordinated functional gene modules rather than individual gene changes. Despite the various types of nanoparticles, prolonged exposure stimulated a common molecular mechanism, including the induction of mitochondrial ISR, antioxidant response, mitochondrial adaptation, and inflammatory signaling, while maintaining specific features of nanoparticle gene expression.

**Figure 5.**
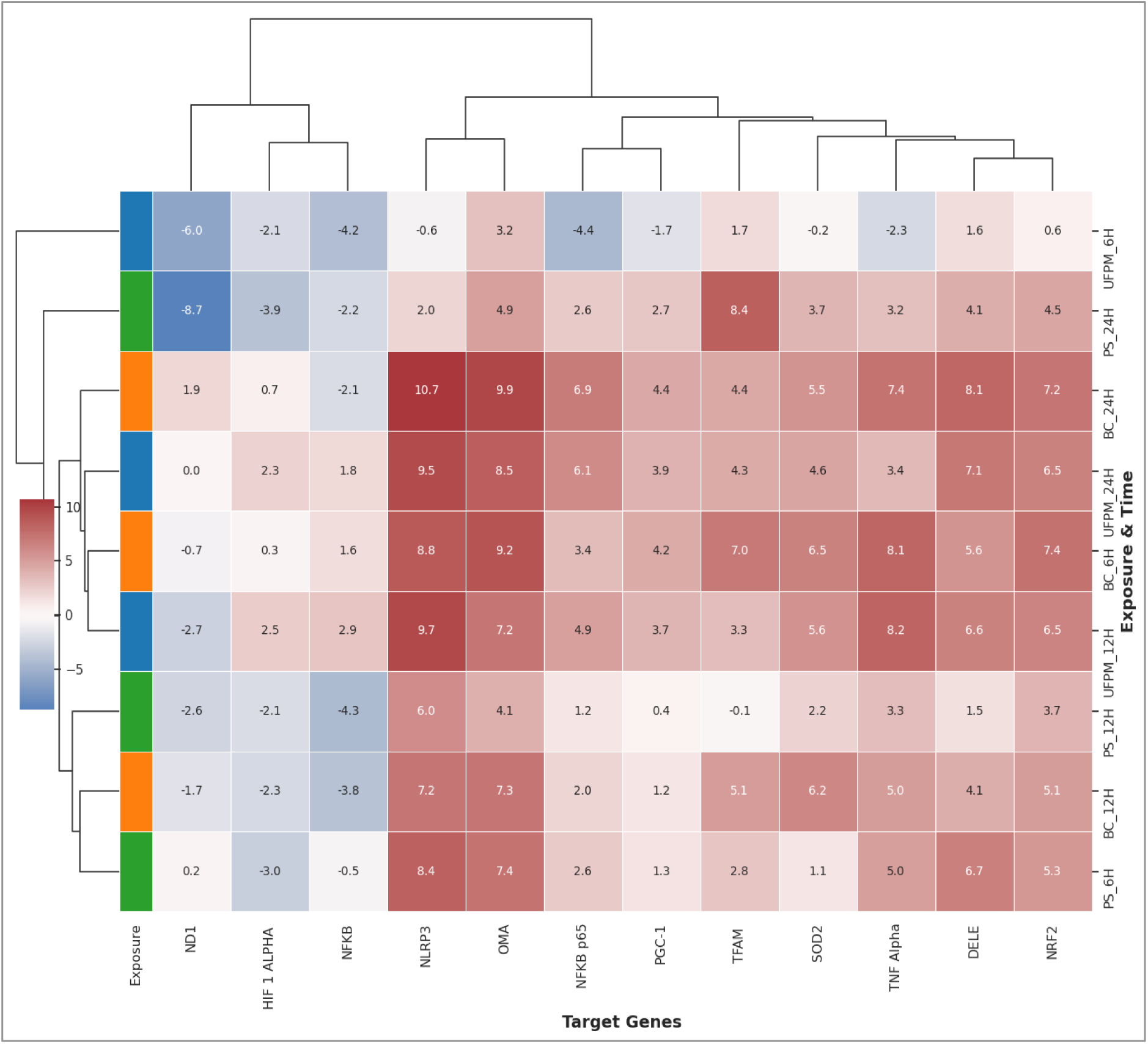
Hierarchical clustering of average gene expression profiles upon exposure to BC, PS-NPs, and UFPM. Heat map showing hierarchical clustering of average log₂ fold change values of genes involved in mitochondrial stress, inflammation, oxidative stress, hypoxia, and mitochondrial biogenesis in response to BC, PS-NPs, and UFPM at 6, 12, and 24 h. Rows are exposure time combinations, while columns are the target genes (ND1, HIF-1α, NF-κB, NLRP3, OMA1, NF-κB p65, PGC-1α, TFAM, SOD2, TNF-α, DELE1, and NRF2). Hierarchical clustering analysis was carried out using Euclidean distance and Ward’s linkage algorithm to detect similarities in exposure and gene expression profile. Colors in cells represent average log₂ fold change values, with blue representing relatively lower expression and red representing relatively higher expression. The number in each cell is the corresponding value of expression.

### 3.6. Gene-to-gene correlation analysis reveals coordinated mitochondrial stress networks

The Pearson correlation analysis was used to investigate the interaction of genes responsible for mitochondrial stress, oxidative stress, mitochondrial bioenergetics, and inflammatory signaling under nanoparticle exposure (Figure 6). Generally, the correlation matrix showed mostly strong and moderate correlations between the analyzed genes, indicating their simultaneous regulation at the level of transcription rather than individual pathway activity. There were no negative correlations, meaning that mitochondrial stress response, antioxidant protection, mitochondrial adaptation, and inflammatory responses were activated in a highly integrated way. Strong correlations were shown between the genes involved in different biological pathways. The highest correlations were found between TNF-α and NRF2 (r = 0.86), PGC-1α and SOD2 (r = 0.84), DELE1 and NRF2 (r = 0.84), NRF2 and NF-κB p65 (r = 0.83), DELE1 and NF-κB p65 (r = 0.82), and ND1 and OMA1 (r = 0.80). Moreover, there were some strong correlations between OMA1, PGC-1α, NLRP3, TNF-α, SOD2, and NRF2. These results showed that there is a great deal of interaction between the pathways of mitochondrial ISR, antioxidant response, mitochondrial biogenesis, and inflammation.

**Figure 6.**
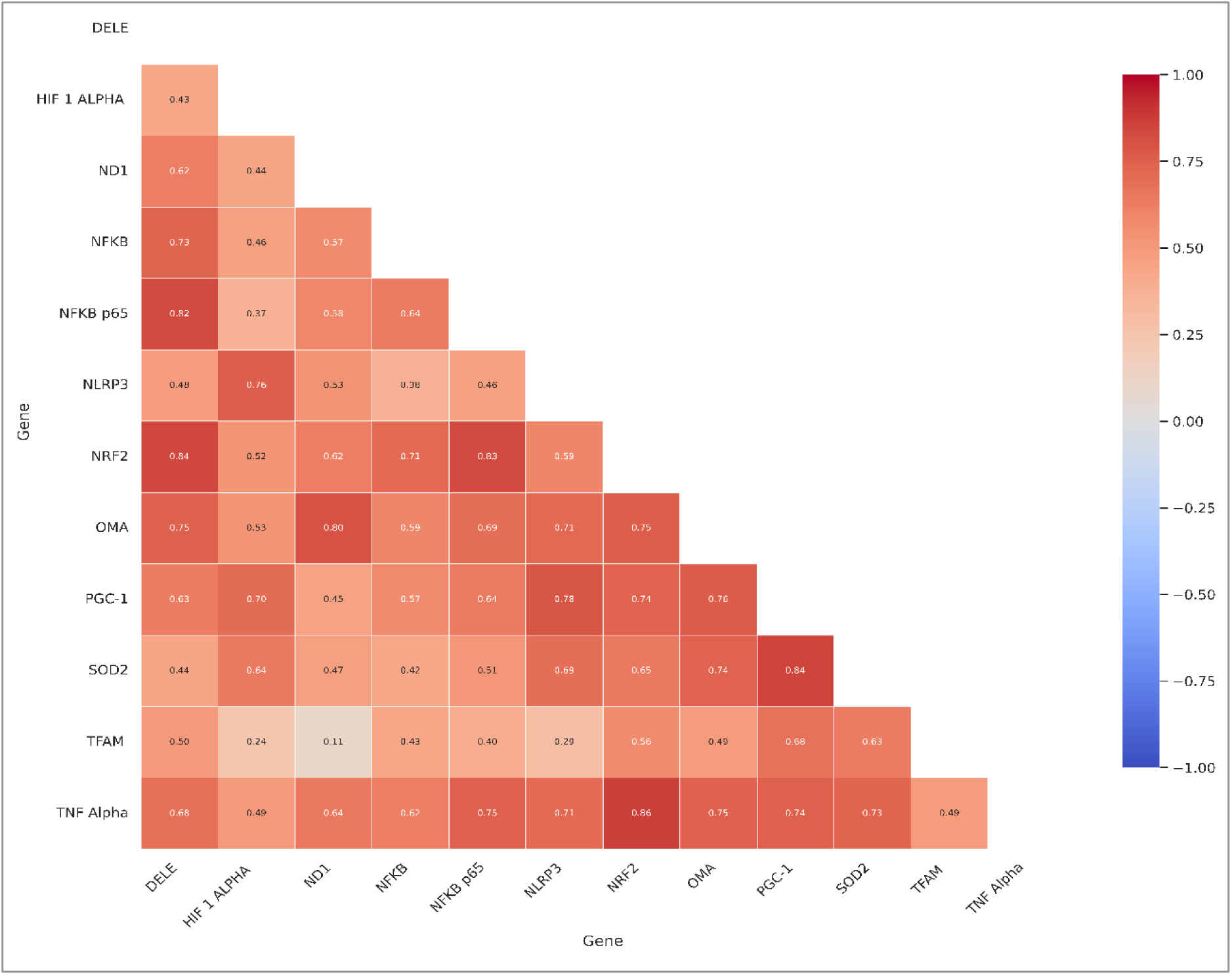
Pearson correlation between gene expressions on mitochondrial stress, inflammatory, oxidative stress, and mitochondrial biogenesis pathways. Heat map showing Pearson correlation coefficients (r) between the expression levels of DELE1, HIF-1α, ND1, NF-κB, NF-κB p65, NLRP3, NRF2, OMA1, PGC-1α, SOD2, TFAM, and TNF-α at all exposure conditions and different time points. The lower triangular matrix shows the Pearson correlation coefficient between all gene pairs, with the value of r in each box. Color intensity indicates the degree and direction of correlation, ranging from −1 (strong negative correlation; blue) to +1 (strong positive correlation; red), as shown in the color scale. Pearson correlation was used to assess transcriptional coordination among genes related to mitochondrial quality control, oxidative stress response, inflammatory signaling, hypoxia, and mitochondrial biogenesis after exposure to BC, PS, and UFPM.

### 3.7. Shared variance analysis supports coordinated regulation of mitochondrial stress pathways

In addition, the coefficient of determination (R²) was obtained to estimate the amount of shared variance between the gene expression profiles and evaluate the coordinated regulation of mitochondrial stress response pathways (Figure 7). Like the results from the Pearson correlation analysis, some genes responsible for mitochondrial stress, oxidative stress, and inflammation displayed considerable shared variance, implying that a large portion of their transcriptional variation could be explained by their shared regulatory responses. The genes with the greatest shared variance included TNF-α and NRF2 (R² = 0.74), PGC-1α and SOD2 (R² = 0.71), DELE1 and NRF2 (R² = 0.70), NRF2 and NF-κB p65 (R² = 0.68), DELE1 and NF-κB p65 (R² = 0.67), as well as ND1 and OMA1 (R² = 0.64). However, high common variance between OMA1, PGC-1α, NLRP3, TNF-α, and SOD2 suggests correlations among mitochondrial ISR regulation, antioxidant, bioenergetic, and inflammatory signaling pathways. On the other hand, the TFAM gene showed low common variance with other genes such as ND1 (R² = 0.01), HIF-1α (R² = 0.06), and NLRP3 (R² = 0.09). Overall, the analysis of shared variance confirmed the presence of a well-integrated network of mitochondrial stress response pathways, although it also showed the independence of certain pathway components, such as TFAM, from other pathways under nanoparticle-induced cell stress.

**Figure 7.**
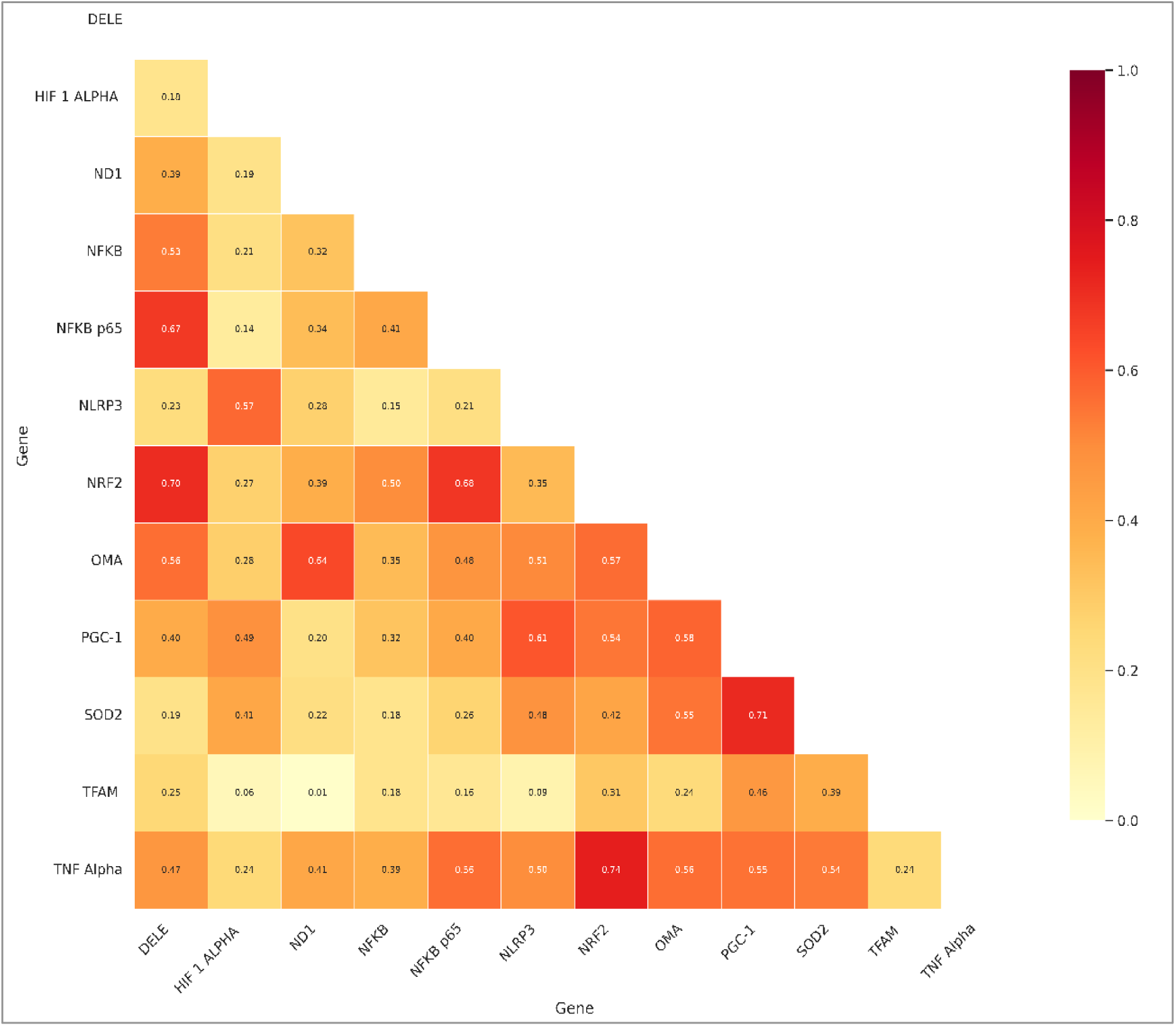
Shared variance (R²) among genes related to mitochondrial stress, inflammatory signaling, oxidative stress, and mitochondrial biogenesis. Heat map showing the pairwise coefficients of determination (R2) obtained using Pearson correlation analysis of the expression patterns of DELE1, HIF-1α, ND1, NF-κB, NF-κB p65, NLRP3, NRF2, OMA1, PGC-1α, SOD2, TFAM, and TNF-α at all exposure times and doses. R² values for each gene pair are shown in the lower triangle of the matrix and reflect the amount of shared variance in the gene expression profile. Numbers within each square show the R² values, while the color scale shows the extent of shared variance, where the light-yellow color represents 0 (low level of shared variance), whereas the dark red color indicates 1 (high level of shared variance). The objective of this analysis was to identify the extent of common transcriptional variability in genes associated with mitochondrial quality control, hypoxia, oxidative stress, inflammation, and mitochondrial biogenesis following exposure to BC, PS-NPs, and UFPM.

### 3.8. Pairwise significance analysis validates coordinated mitochondrial stress networks

Statistical significance analysis was carried out to assess the strength of gene-gene relationships established using correlation and variance analyses (Figure 8). As shown in the Pearson correlation matrix, most interactions among genes involved in mitochondrial stress, oxidative stress, bioenergetics, and inflammatory pathways were statistically significant (p < 0.05). More specifically, some interactions had an extremely high level of statistical significance (p < 0.001) and included such genes as DELE1, OMA1, NRF2, PGC-1α, SOD2, TNF-α, NF-κB, and NF-κB p65. All these findings confirm the coordinated regulation of mitochondrial ISR, antioxidant defense, mitochondrial adaptations, and inflammatory pathways under conditions of nanoparticle exposure. At the same time, some genes, including TFAM, showed fewer significant interactions with other genes; namely, HIF-1α, ND1, and NLRP3 were not significantly associated with TFAM, and the interaction between HIF-1α and NF-κB p65 was not statistically significant. These data suggest that although mitochondrial biogenesis is part of the general stress response network, some genes responsible for mitochondrial genome integrity demonstrate greater transcriptional independence than genes responsible for mitochondrial ISR, oxidative stress, and inflammatory signaling. Overall, the significance analysis confirms the validity of the stress response network established using clustering and correlation analyses.

**Figure 8.**
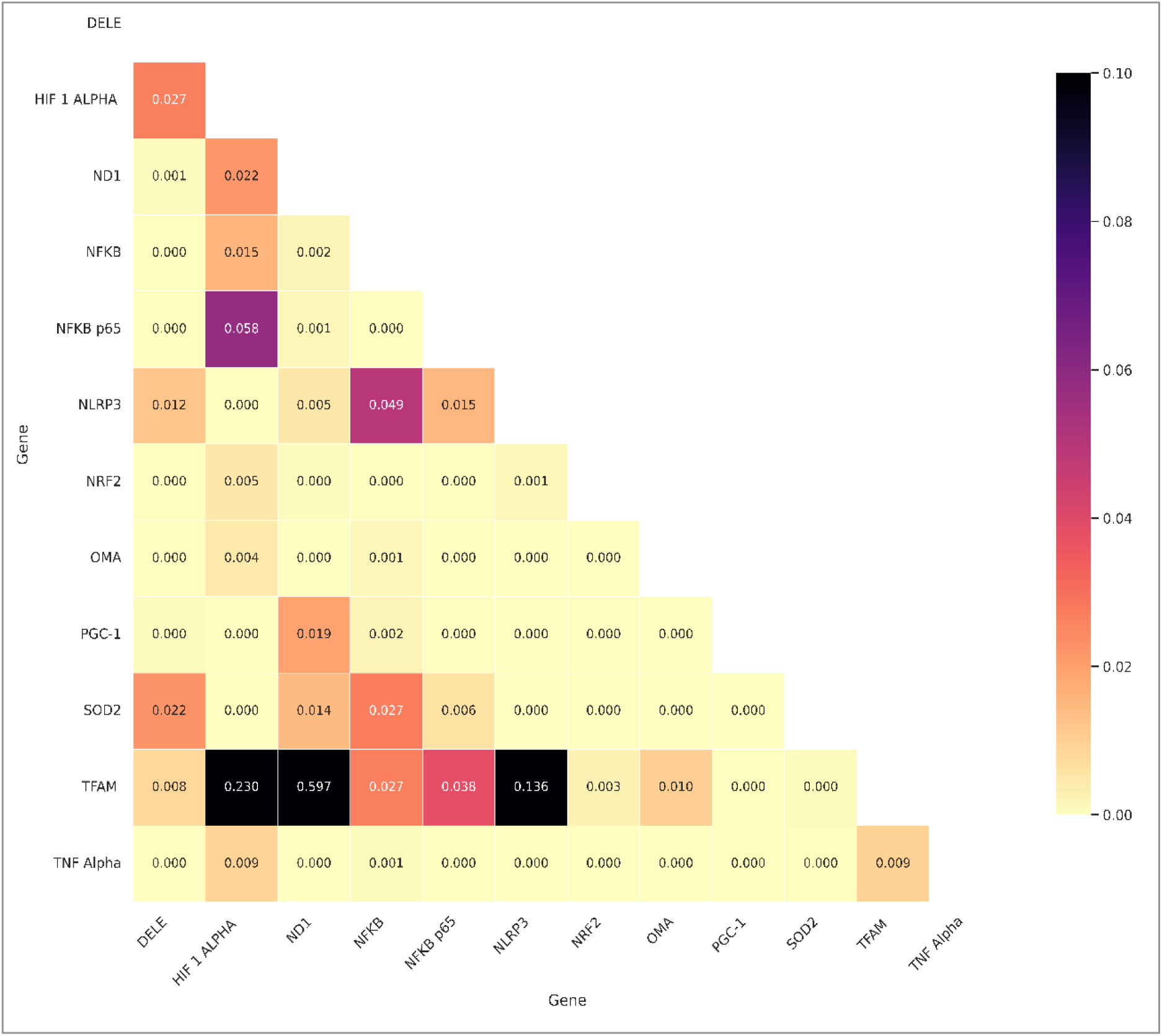
Statistical significance of pairwise gene-gene correlations in mitochondrial stress, inflammation, oxidative stress, and mitochondrial biogenesis pathways. Heat map indicating the p-values resulting from the pairwise Pearson correlation analysis on the expression profiles of DELE1, HIF-1α, ND1, NF-κB, NF-κB p65, NLRP3, NRF2, OMA1, PGC-1α, SOD2, TFAM, and TNF-α under all exposure conditions and all time points. The lower triangular portion of the matrix indicates the p-value for each gene pair, with the numbers presented within each cell. Color intensity denotes the significance of the p-values, with lighter colors indicating low p-values (high statistical significance), and vice versa, according to the color scale provided. In this study, the statistical significance of interactions involving transcriptional regulation of genes involved in mitochondrial quality control, hypoxia signaling, oxidative stress, inflammation, and mitochondrial biogenesis following BC, PS, and UFPM treatment was analyzed.

### 3.9. Multi-omics correlation links mitochondrial stress pathways with inflammatory cytokine secretion

To test for a possible downstream link between transcriptional alterations and inflammation, Pearson’s correlation analysis was applied to compare the expression levels of mitochondrial stress-related genes with cytokine concentrations detected by ELISA (Figure 9). Cytokine-specific correlation patterns have been revealed. IL-6 and IL-8 showed high correlation with the expression levels of genes related to mitochondrial stress, oxidative stress, and inflammation, whereas IL-1β showed low or even negative correlations. TNF-α secretion showed moderate correlations with several mitochondrial stress-induced genes. Among the cytokines considered, IL-6 has the strongest correlations with mitochondrial stress pathways, including high positive correlations with HIF-1α and NF-κB (r = 0.73), together with moderate correlations with DELE1, NLRP3, OMA1, PGC-1α, NRF2, and ND1. Similarly, IL-8 had strong correlations with genes participating in mitochondrial ISR, oxidative stress response, and inflammation, including high positive correlations with NF-κB p65 (r = 0.72), PGC-1α (r = 0.58), SOD2 (r = 0.52), NLRP3 (r = 0.49), and NRF2 (r = 0.49). In contrast, TNF-α release was characterized by moderate positive correlations with ND1, HIF-1α, Mitogen ND1, NLRP3, OMA1, and SOD2, whereas IL-1β revealed only weak correlations with most of the genes and negative correlations with several mitochondria-specific proteins, including especially TFAM (r = -0.48). Altogether, the multi-omics data integration indicates the close relationship between transcriptional activation of mitochondrial stress pathways and cytokine secretion upon nanoparticle treatment. Genes responsible for mitochondrial ISR (OMA1-DELE1), oxidative stress response (NRF2-SOD2), hypoxic adaptation (HIF-1α), bioenergetics (ND1), and inflammatory signaling (NF-κB/NF-κB p65 and NLRP3) were selectively correlated with IL-6 and IL-8, which can be considered as the main outputs of inflammatory responses related to mitochondrial stress. Conversely, the relatively weak correlations of TFAM with all analyzed cytokines, especially IL-1β and TNF-α, imply independent regulation of mitochondrial biogenesis and DNA maintenance from acute cytokine secretion. The presented results provide integrated transcriptomic-proteomic evidence about the coupling of mitochondrial dysfunction with downstream inflammation in response to environmentally relevant nanoparticles with distinct chemical composition.

**Figure 9.**
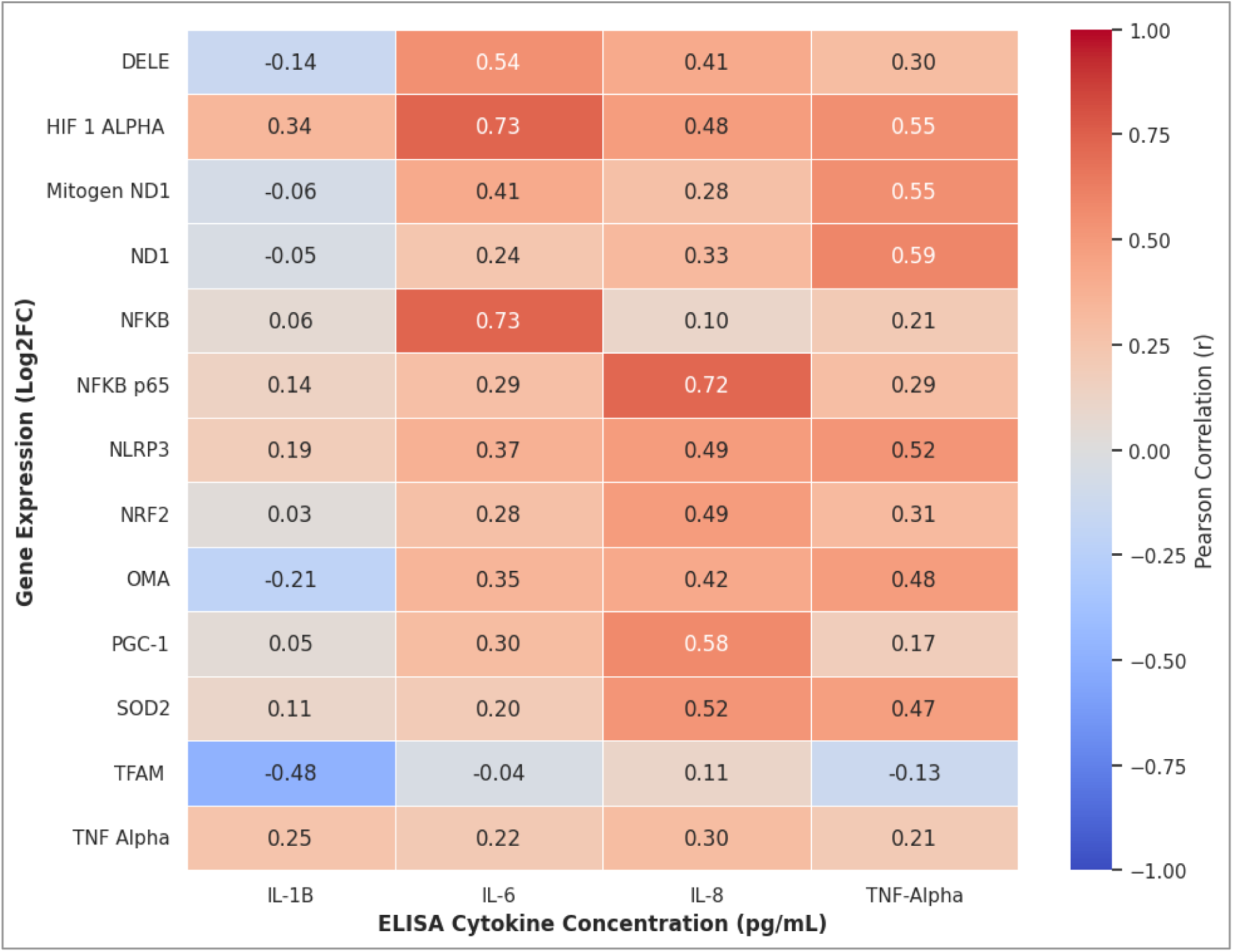
Correlation of gene transcription and cytokine secretion in response to airborne particulate pollutants. Heat map depicting Pearson correlation coefficients (r) of the relative expression of genes related to mitochondrial stress, inflammation, oxidative stress, and mitochondrial biogenesis and the level of cytokines measured using ELISA. Log2-transformed fold changes in gene expression for DELE1, HIF-1α, Mitogen ND1, ND1, NF-κB, NF-κB p65, NLRP3, NRF2, OMA1, PGC-1α, SOD2, TFAM, and TNF-α were correlated with the levels of IL-1β, IL-6, IL-8, and TNF-α at all exposure conditions and time points. The numeric value in each cell represents the Pearson correlation coefficient (r), while the color scale indicates the degree of correlation, which can vary from -1 (negative correlation; blue) to +1 (positive correlation; red). This holistic approach investigated the correlation between transcription and cytokine secretion in response to BC, PS-NPs, and UFPM.

Collectively, these transcriptomic, statistical, and multi-omics studies provide evidence for the induction of a conserved yet dynamically regulated mitochondrial stress response by chemically different environmental nanoparticles. Although there were differential gene expression levels and dynamics, UFPM, BC, and PS-NPs activated a similar pattern of pathways, including mitochondrial ISR (OMA1-DELE1), oxidative stress (NRF2-SOD2), mitochondrial metabolic adaptation (HIF-1α-PGC-1α-TFAM-ND1), and inflammation (NF-κB, NLRP3, and TNF-α). Using systems biology approaches such as PCA, hierarchical clustering, correlation, shared variance, and transcript and cytokine associations, we formed a clear picture of a highly connected network between mitochondrial function and cytokine secretion. Whereas BC caused sustained mitochondrial stress and an inflammatory response, UFPM caused the most dynamic transcriptional reprogramming, and PS-NPs were more inclined to mitochondrial adaptation to an inflammatory response. Taken together, these results suggest a common mitochondrial stress adaptation mechanism as the key driver of the immune toxicity of environmental nanoparticles with specific particle-dependent molecular signatures.

## 4. Discussion

As revealed in the current study, diverse environmental nanoparticles exhibit a similar mitochondrial stress response despite the high chemical diversity. UFPM, BC, and PS-NPs were studied for their effects on human PBMCs, showing similar downstream responses associated with oxidative stress, energy production, ISR activation, and inflammation. Although separate studies reported negative effects of UFPMs, BC, and PS-NPs on mitochondria (Mishra et al., 2022; Shang et al., 2022; Lee et al., 2022), comparative studies are lacking. While all three nanoparticles showed activation of overlapping stress markers, there were clear differences in the magnitude and the time course of these responses depending on the chemical nature of the particles, with rapid adaptation caused by UFPM, chronic mitochondrial dysfunction and inflammatory activation caused by BC, and relatively moderate bioenergetic remodeling caused by PS-NPs. Our results provide further evidence for the emerging idea that mitochondria are key environmental stress sensors capable of integrating oxidative, metabolic, and inflammatory signals into a coordinated stress response rather than responding separately to different pollutants (Quirós et al., 2016; Guo et al., 2020; Fessler et al., 2020).

Our results imply that mitochondrial redox dysregulation is probably one of the critical initiating events after UFPM and BC exposures, whereas PS-NPs might have slightly different pathways leading to the same result. Redox-sensitive genes were all upregulated in response to NP exposure, though with different kinetics depending on the nature of the NP. While UFPM caused a quick and transient response of the antioxidant defense system, this is likely due to a high level of reactivity of its surface and the presence of transition metal ions and quinones, which facilitate redox cycling and Fenton-type reactions (Li et al., 2003; Xia et al., 2006; Nel et al., 2006; Bhargava et al., 2017). In contrast, BC was linked to continuous expression of the NRF2 gene, implying oxidative stress signaling due to continuous ROS production from combustion products such as polycyclic aromatic hydrocarbons, among others. This study confirmed previous literature that BC treatment causes chronic oxidative stress, which results in mitochondrial-specific oxidative stress (Janssen et al., 2011; Sharma et al., 2020; Shang et al., 2022; Schraufnagel, 2020). Lastly, PS-NPs induce mitochondrial stress through their interactions inside cells rather than their surface reactions (Cedervall et al., 2007; Monopoli et al., 2012; Lee et al., 2022). Strong positive correlations between NRF2-TNF-α (r = 0.86), NRF2-DELE1 (r = 0.84), and PGC-1α-SOD2 (r = 0.84) are consistent with mitochondrial oxidative stress being associated with mitochondrial quality control and adaptation responses. Remarkably, PS-NPs induced downstream effects related to mitochondrial stress in the absence of a strong oxidative stress response, implying that mitochondrial disturbances could also be triggered in the absence of oxidative stress through other pathways, including changes in mitochondrial dynamics and interactions. This shows that oxidative stress is just one of many possible triggers of a mitochondrial stress response.

Apart from inducing oxidative stress, our results suggest that environmental nanoparticles are associated with transcriptional signatures consistent with bioenergetic remodeling rather than merely affecting their function. Concomitant changes in HIF-1α, PGC-1α, TFAM, ND1, and Complex I-V activity are consistent with a dynamic transcriptional response to oxidative stress for the maintenance of energy balance in cells (Scarpulla, 2008; Scarpulla, 2011; Spinelli & Haigis, 2018). The intensity of such remodeling, nevertheless, differed among nanoparticle types. UFPM caused potent HIF-1α activation along with the normalization of PGC-1α and an increase in TFAM, indicating efficient metabolic and mitochondrial adaptations after oxidative stress (Semenza, 2012; Chandel, 2015). In BC, HIF-1α is activated, and the respiratory rate is low, suggesting that mitochondrial dysfunction in BC does not serve only an adaptive purpose (Murphy, 2009; Bhargava et al., 2017; Sharma et al., 2020; Shang et al., 2022). Non-provocative PS-NPs increased the expression of TFAM, which agrees with previous results concerning the impact of nanoplastics on mitochondrial DNA and mitochondrial dynamics before cell damage (Lee et al., 2022). The correlation between ND1 and OMA1 (r = 0.80), as well as between PGC-1α and SOD2 (r = 0.84), indicates that respiration, mitochondrial production, and antioxidants are all linked in a coordinated mechanism. In consequence, although all three nanoparticle types evoke a similar mitochondrial stress response, the chemical nature of the particles determines whether mitochondrial bioenergetic adaptation occurs successfully (Mishra et al., 2022). Yet another important finding is the coordinated activation of OMA1 and DELE1 among all three nanoparticle types, showing that mitochondrial ISR is an evolutionarily conserved mechanism connecting mitochondrial stress with cellular adaptive stress response. The OMA1-DELE1 pathway has recently gained recognition as an essential player in regulating mitochondrial protein quality control, although its role in nanoparticle-induced mitochondrial stress is poorly studied (Guo et al., 2020; Fessler et al., 2020; Mick et al., 2020). Despite the great dissimilarity in oxidative stress kinetics, Treatment with UFPM, BC, and PS-NPs was associated with increased OMA1 and DELE1 transcript expression, consistent with engagement of the mitochondrial ISR pathway. BC caused the strongest sustained induction of OMA1 and DELE1 due to persistent oxidative stress and impaired respiration; however, the response to UFPM was much more transient, and the activation of the molecular pathway in PS-NPs was less pronounced, meaning better mitigation of mitochondrial dysfunction. In addition, the high positive correlation obtained between DELE1 and NRF2 (r = 0.84), as well as that between ND1 and OMA1 (r = 0.80), suggests a strong relationship between oxidative stress, respiratory distress, and mitochondrial surveillance. These results add to current literature and demonstrate the involvement of the OMA1-DELE1-mitochondrial ISR pathway in nanoparticle-triggered mitochondrial adaptation, an exciting novel mechanism and marker of nanoparticle-induced mitochondrial stress (Mishra et al., 2022; Wang & Zhang, 2024).

Further, our results indicate that inflammation activation is a downstream effect of mitochondrial stress activation rather than an independent effect caused by exposure to nanoparticles. In accordance with our proposed model, NF-κB, NF-κB p65, TNF-α, and NLRP3 are primarily expressed during later time exposure phases. Therefore, activation of inflammation was secondary to the initial response to bioenergetics and oxidative stress. Of the three categories of nanoparticles studied, BC resulted in the greatest inflammatory activity, as indicated by the up-regulation of genes that are products of NF-κB and NLRP3. UFPM had relatively transitory responses, and PS-NPs induced moderate inflammation in the absence of substantial mitochondrial reprogramming. These results are consistent with previous studies reporting that inflammation and epigenetic effects occur due to NF-κB activation after UFPM treatment (Bhargava et al., 2019).

In contrast to previous studies comparing one particle class at a time, our current study provides evidence that all three nanoparticles activate a mitochondrial stress response program in the same experimental setting (West et al., 2015; Riley & Tait, 2020; Shang et al., 2022). Furthermore, the simultaneous stimulation of NLRP3, NF-κB, and TNF-α signaling pathways implies the priming function of the inflammasome mediated by the danger signal emitted by mitochondria, consistent with past research demonstrating the importance of ultrafine particles and nanoplastics in NLRP3 activation via the mitochondrial oxidative stress pathway (Zhou et al., 2011; Swanson et al., 2019; Zou et al., 2026; Lee et al., 2022). In addition, the strong correlation between TNF-α and NRF2 (r = 0.86), as well as between HIF-1α and NLRP3 (r = 0.76), demonstrates a close connection between inflammation, oxidative stress, and metabolic alterations, instead of independent reactions. In general, the data discussed above confirm the model whereby chronic mitochondrial stress shifts the body toward inflammation.

The multivariate statistical analysis independently validates the mechanism-driven model outlined in this study, demonstrating that nanoparticle-induced mitochondrial stress is a consequence of biological processes occurring in unison rather than individually. The proportion of variance described by the first two principal components (PC1 and PC2) using PCA was 73.1% (PC1 - 60.9%, PC2 - 12.2%). This means that most of the variability is attributed to the regulation of oxidative stress, mitochondrial remodeling, and the inflammatory response. Based on these molecular findings, the UFPM cluster showed the highest degree of temporal variability, indicating that mitochondrial adaptation is dynamic. On the other hand, BCs mostly accumulated along the positive PC1 axis, which means that mitochondria and inflammation remain constantly dysregulated. The location of PS-NPs is quite variable, indicating some mitochondrial adaptation despite low inflammation. Moreover, hierarchical clustering showed that exposure duration has a higher impact on transcriptional organization than particle type, corroborating the idea that different chemically distinct nanoparticles become more similar in their effects with prolonged exposure through induction of a common mitochondrial stress phenotype. Both correlation and regression analyses support this hypothesis, as there is a high correlation among NRF2, DELE1, PGC-1α, SOD2, ND1, OMA1, TNF-α, and NLRP3. This implies that there is a well-coordinated interaction between oxidative stress, mitochondrial quality control, metabolic remodeling, and inflammation. Therefore, system biology studies show the independence of statistical proof, showing that physical and chemical properties of particles influence the kinetics of mitochondrial adaptation but not the stress response pathways (Kitano, 2002; Barabási et al., 2011; Hasin et al., 2017; Ringnér, 2008; Lever et al., 2017; Mishra et al., 2022).

The results of our study not only support findings from other studies on how UFPM, BC, and PS-NPs can cause mitochondrial dysfunction but also build on existing knowledge (Byun et al., 2013; Shang et al., 2022; Lee et al., 2022). Contrary to prior work, which has studied mitochondria after exposure to individual nanoparticle classes in a variety of experimental models, this study enables direct comparison of these particles under the same experimental conditions to investigate both common and particle-specific effects on the mitochondrial system. It becomes apparent that a wide range of nanoparticles induces a convergent response at the level of mitochondrial stress response pathways but shows differences only in terms of the intensity and dynamics of mitochondrial adaptation, thus confirming and expanding upon the previously observed mitochondrial dysfunction and mitoepigenetic regulation caused by the exposure to particulate matter (Bhargava et al., 2017; Sharma et al., 2020; Mishra et al., 2022). Human PBMCs used in the current study increase the translational importance of the findings, since these circulating immune cells are a readily available proxy for systemic exposure and can be effectively employed as biomonitoring tools in environmental monitoring efforts (Hou et al., 2013; Janssen et al., 2011). However, several issues should be addressed. Acute ex vivo exposure does not fully recapitulate chronic environmental conditions, particle accumulation in tissues, or organ-specific responses. In addition, functional validation of the signaling pathway and mitochondrial bioenergetics is not included in the current study. Further research combining in vivo exposure models and approaches such as single-cell transcriptomics, spatial transcriptomics, phosphoproteomics, and mitochondrial metabolomics will be needed to ensure physiological relevance of the proposed pathway (Hasin et al., 2017; Stuart & Satija, 2019; Marx, 2021). From an applied point of view, the regulatory coordination among OMA1, DELE1, NRF2, PGC-1α, TFAM, and ND1 indicates that these genes associated with the mitochondrial ISR may be potential biomarkers that need to be validated in subsequent studies (Kelly & Fussell, 2015; Schraufnagel, 2020). In general, these results reveal mitochondrial stress adaptation as a common mechanism underpinning nanoparticle toxicity and pave the way toward developing mechanism-based biomarkers of environmental exposure.

## 5. Conclusion

The current study illustrates that chemically different environmental nanoparticles (UFPM, BC, and PS-NPs) share a conserved mitochondrial stress response program despite being physically different from one another. In fact, by directly comparing these relevant environmental nanoparticles under identical conditions in human PBMCs, we provide evidence that oxidative stress, mitochondrial bioenergetic reprogramming, mitochondrial ISR through the OMA1-DELE1 pathway, and inflammation comprise a network of interrelated responses, the extent and kinetics of which depend mostly on the chemical composition of the nanoparticle. Thus, our work suggests that mitochondrial dysfunction is a dynamic process regulated by coordinated redox homeostasis, mitochondrial quality control, metabolic reprogramming, and activation of innate immunity rather than a simple toxicological effect. In addition, comparative systems biology analyses of these three diverse nanoparticles have shown that the OMA1-DELE1-dependent ISR is an important molecular link between oxidative stress and subsequent inflammation. Additionally, we revealed that a complex network of biomarkers comprising NRF2, PGC-1α, TFAM, ND1, OMA1, DELE1, TNF-α, and NLRP3, as well as mitochondrial respiratory complexes I-V, is functionally integrated. Altogether, we demonstrate that mitochondrial stress adaptation is a unified mechanistic basis of the toxicity of environmental nanoparticles with different chemical compositions, advance our understanding of particle-specific mitochondrial responses, and lay a strong foundation for the development of biomarker panels and mitochondria-targeted therapeutic strategies for environmental risk assessment and precision toxicology.

## Abbreviations

BC: Black carbon
DELE1: DAP3-binding cell death enhancer 1
HIF-1α: Hypoxia-inducible factor-1 alpha
ISR: Integrated stress response
mtDNA: Mitochondrial DNA
ND1: NADH dehydrogenase subunit 1
NF-κB: Nuclear factor kappa B
NLRP3: NOD-like receptor family pyrin domain-containing 3
NRF2: Nuclear factor erythroid 2-related factor 2
OMA1: Overlapping activity with m-AAA protease 1
PBMCs: Peripheral blood mononuclear cells
PGC-1α: Peroxisome proliferator-activated receptor gamma coactivator-1 alpha
PS-NPs: Polystyrene nanoplastics
ROS: Reactive oxygen species
TFAM: Mitochondrial transcription factor A
TNF-α: Tumor necrosis factor-alpha
UFPM: Ultrafine particulate matter;
Δψm: Mitochondrial membrane potential.

## Declaration

### Ethics approval and consent to participate

The study received approval from the Institutional Ethics Committee (IEC) of ICMR-NIREH under reference number INTR-IM-2024-00063 **(PKM).**

### Consent for publication

Not applicable

### Availability of data and materials

Data will be made available on request.

### Authors’ contributions

PKM conceptualized the study, developed the methodology, provided supervision, managed project administration, and secured funding acquisition. AC, AKR, AP, AA, PPD, and RPT performed the investigation and executed the experiments. VG and AA were responsible for sample collection and physicochemical characterization of the materials. VG and DD handled data visualization and figure design. DKS, RT, and RKS contributed to data analysis and manuscript editing. DD, AKR, AC, VG, and PKM prepared the original draft of the manuscript. All authors reviewed and approved the final version.

## Acknowledgments

The authors are thankful to Mr. Sagar Patel and Mr. Ashish Kumar for their technical assistance.

