## Supplementary methods for "Comparative Analysis of Ultrafine Particulate Matter, Black Carbon, and Polystyrene Nanoplastics Identifies Mitochondrial Stress Adaptation as a Conserved Mechanism of Immunotoxicity"

**2.1 Sample Collection and Lymphocyte Isolation**

Blood samples from healthy adults were drawn via venipuncture with approval from the Institutional Ethics Committee of the ICMR-National Institute for Research in Environmental Health. Blood sampling procedures followed the ethical guidelines. The samples were transported to the laboratory at 2-8 °C within 2-3 h of collection. PBMCs were separated by density gradient centrifugation technique using HiSep™ LSM 1077 (HiMedia Laboratories Pvt. Ltd., Mumbai, India) as per the manufacturer's protocol. The PBMCs were first washed twice with sterile PBS solution and then were pelleted by centrifugation. PBMCs were cultured in the presence of UFPM, BC, or PS-NPs. PBMCs were harvested after 6, 12, and 24 h. PBMCs that were not subjected to any treatment and maintained under similar culture conditions served as controls. The cell pellets were collected after each time point and preserved at -80 °C for further analysis (Bhargava et al., 2017).

**2.2 Treatment and Exposure of UFPM, BC, and PS**

UFPM, BC, and PS stock suspensions were prepared in 1X PBS. Dilutions were prepared to achieve a final concentration of 1 µg/mL. The same dose was applied to all three particulates to facilitate direct comparison under similar conditions. This dose was chosen to study early oxidative stress, mitochondrial dysfunction, and inflammation without causing unnecessary cell death. The cells were exposed to the particulate matter for 6, 12, and 24 hours under a humidified environment of 5% CO2. The control group included unexposed cells. After exposure to each particulate for the respective time duration, the cells were pelleted by centrifugation at 4,500 × g for 10 min. The cells were then washed twice in 1X PBS to remove any unbound particles. Cells were frozen at -80 °C for further processing (Pan et al., 2018; Salinas et al., 2020; Adamiak et al., 2025).

**2.3 Analysis of Oxidative Stress and Apoptosis**

Genomic DNA was isolated from control and treated PBMCs (UFPM, BC, and PS) using the PureLink™ Genomic DNA Mini Kit (Invitrogen, Thermo Fisher Scientific, USA) as per the kit instructions. The oxidation of DNA was determined by measuring oxidized purine bases, such as 8-oxodeoxyguanosine, through an Fpg digestion assay, as described in previous literature (Mishra et al., 2022; Bunkar et al., 2020). The ROS levels inside the cells were measured during assessment of oxidative stress using CM-H₂DCFDA staining. Apoptosis and necrosis were studied using CellEvent™ Caspase-3/7 Green reagent along with SYTOX™ AADvanced™ dead cell stain (Thermo Fisher Scientific, USA). The stained cells were analyzed by flow cytometry using 488 nm excitation, with emission collected at 530/30 BP and 690/50 BP filters. All experiments were carried out with appropriate controls and biological replicates to maintain data consistency.

**2.4 Gene Expression Analysis of Oxidative Stress-Related Markers**

RNA isolation was followed by cDNA synthesis. Relative quantitative PCR was conducted using primers for the NRF2 and SOD2 genes. Normalization was performed using a housekeeping gene, and quantification was calculated using the 2^-∆∆CT method. The experiment was conducted in triplicate (Mishra et al., 2022).

**2.5 Assessment of Mitochondrial Functional Markers**

RNA was isolated from the samples and subjected to reverse transcription into cDNA. The qRT-PCR assay was used to assess the expression levels of HIF-1α, PGC-1α, and TFAM genes. Expression levels were normalized against the internal gene, and the relative expression levels were calculated using the 2^-ΔΔCt method. All analyses were done in triplicate.

**2.6 Assessment of Mitochondrial Dysfunction and mtDNA Integrity**

Mitochondrial membrane potential (ΔΨm) was evaluated to assess mitochondrial dysfunction following exposure to UFPM, BC, and PS-NPs. Following treatment, cells were stained with MitoTracker dye according to the manufacturer's protocol. Since MitoTracker specifically accumulates in mitochondria with an intact membrane potential, a decrease in fluorescence intensity was taken as a measure of mitochondrial depolarization. Fluorescence intensity was measured using flow cytometry. Mitochondrial stress signaling was assessed by analyzing the expression of DELE1 and HRI using one-step quantitative real-time PCR.  Relative gene expression was calculated by the 2^-ΔΔCt method after normalization with the housekeeping gene (Bhargava et al., 2017).

**2.7 Evaluation of Mitochondrial Complex Activity Assay**

PBMCs were exposed to UFPM, BC, and PS (1µg/mL) for 6, 12, and 24 h. Cells were collected by centrifugation at 4,500 × g for 10 min. Total protein was extracted from control and treated cells. Mitochondrial electron transport chain (ETC) complex activities were measured using MitoCheck Complex Activity Assay Kits (Cayman Chemical, Ann Arbor, MI, USA) according to the manufacturer's instructions. Absorbance readings were taken using a Spark® multimode microplate reader (Tecan, Männedorf, Switzerland) (Soni et al., 2025a).

**2.8 Immunotoxicity and Inflammatory Response Assessment**

**2.8.1 Cytokine Quantification**

Supernatants from cultures treated with UFPM, BC, and PS-NPs, as well as control cultures, were obtained. They were then centrifuged and stored in the deep freezer at -80°C for future analysis. The concentrations of IL-6, IL-8, and TNF-α in each sample were analyzed using Human GENLISA™ ELISA Kits (KRISHGEN BioSystems, Cerritos, CA, USA). Absorbance of each sample was determined at 450 nm using a Spark® multimode microplate reader (Tecan, Männedorf, Switzerland). The amount of each cytokine was determined using its calibration curve. High cytokine levels in treated samples were interpreted as inflammation (Soni et al., 2025b).

**2.8.2 Assessment of NF-κB Signaling Pathway**

Total RNA from both control and treated PBMCs was isolated using the TRIzol™ reagent (Life Technologies, Thermo Fisher Scientific, Waltham, MA, USA). The total RNA concentration and purity were assessed. The first-strand cDNAs were synthesized from 1 μg of total RNA by using the iScript™ cDNA synthesis kit (Bio-Rad Laboratories, Hercules, CA, USA). The RELA and NFKB1 mRNA expression levels were quantified using the qRT-PCR method. The mRNA expression levels were normalized to an internal reference gene (Mishra et al., 2022).

**2.8.3 Assessment of NLRP3 inflammasome Activation**

Total RNA was extracted from both control and treated PBMCs. RNA concentration and purity were determined spectrophotometrically. cDNA was synthesized from equal amounts of RNA using a reverse transcription kit. The relative expression of the NLRP3 gene was estimated using qPCR with gene-specific primers. The relative expression was normalized to the internal gene using the 2^-ΔΔCt method. IL-1β expression in culture supernatants was estimated by ELISA. Culture supernatants after exposure to particles were centrifuged to remove cell debris and stored at -80 °C for further analysis. IL-1β levels were measured using a human IL-1β ELISA kit according to the manufacturer's protocol. The absorbance was recorded at 450 nm using a Spark® multimode microplate reader (Tecan, Männedorf, Switzerland). The concentration of cytokines was determined from the standard curve (Li et al., 2025).

**2.9 Statistical Analysis**

All computations were done in the programming environment Python. For data analysis, the Pandas and NumPy packages were used. All datasets were standardized using the StandardScaler method of the scikit-learn package prior to performing the analysis. PCA was used for identifying the key factors of variation between the samples. K-means clustering was used to classify the samples based on gene expression patterns. Correlation between the gene expression level and metabolic parameters was estimated using Pearson’s correlation coefficient (Kaur et al., 2025).
