## Supplementary Figs. S1 and S2 for "Comparative Analysis of Ultrafine Particulate Matter, Black Carbon, and Polystyrene Nanoplastics Identifies Mitochondrial Stress Adaptation as a Conserved Mechanism of Immunotoxicity"

### Slide 1
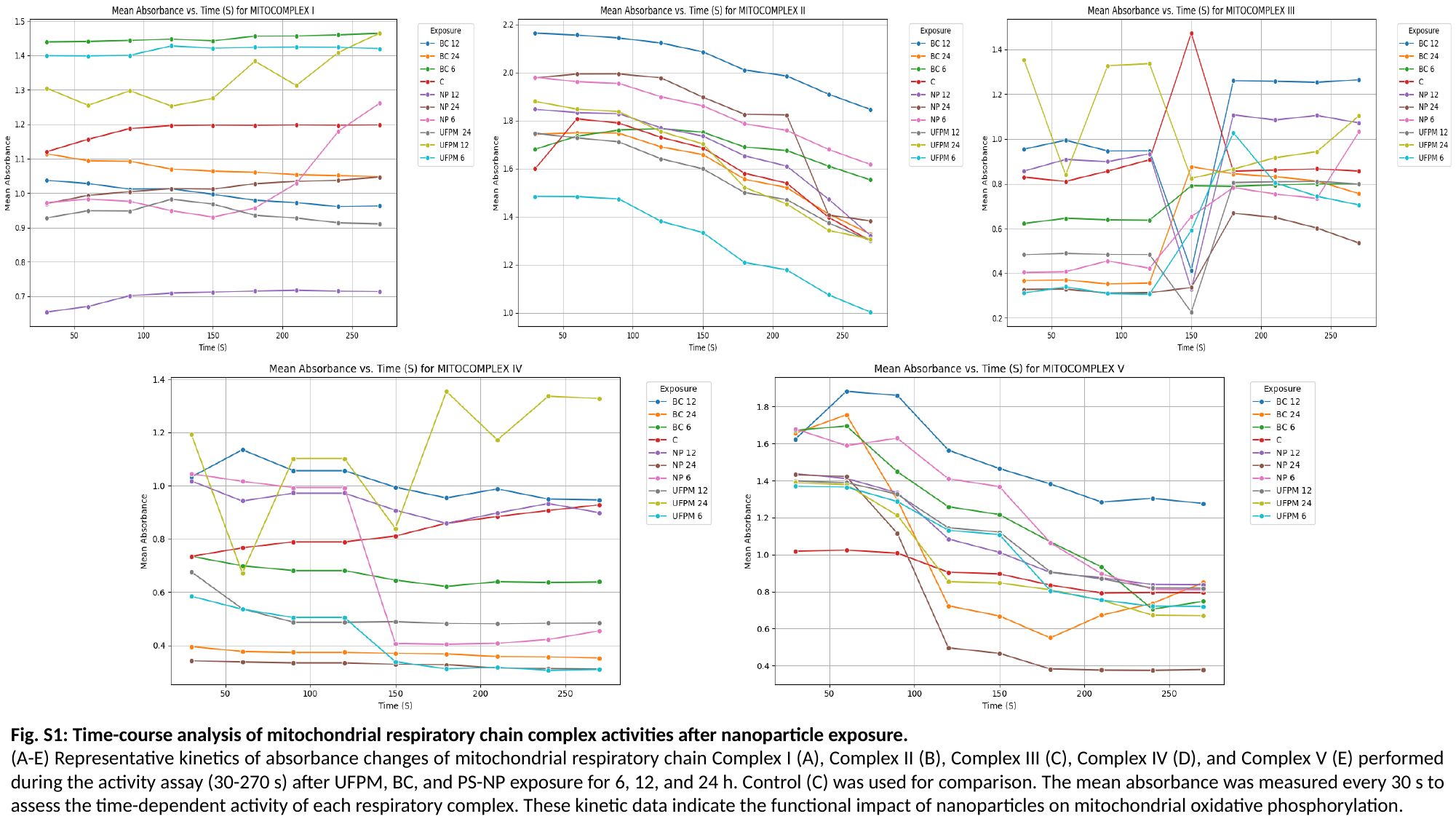

Fig. S1: Time-course analysis of mitochondrial respiratory chain complex activities after nanoparticle exposure.
(A-E) Representative kinetics of absorbance changes of mitochondrial respiratory chain Complex I (A), Complex II (B), Complex III (C), Complex IV (D), and Complex V (E) performed during the activity assay (30-270 s) after UFPM, BC, and PS-NP exposure for 6, 12, and 24 h. Control (C) was used for comparison. The mean absorbance was measured every 30 s to assess the time-dependent activity of each respiratory complex. These kinetic data indicate the functional impact of nanoparticles on mitochondrial oxidative phosphorylation.

### Slide 2
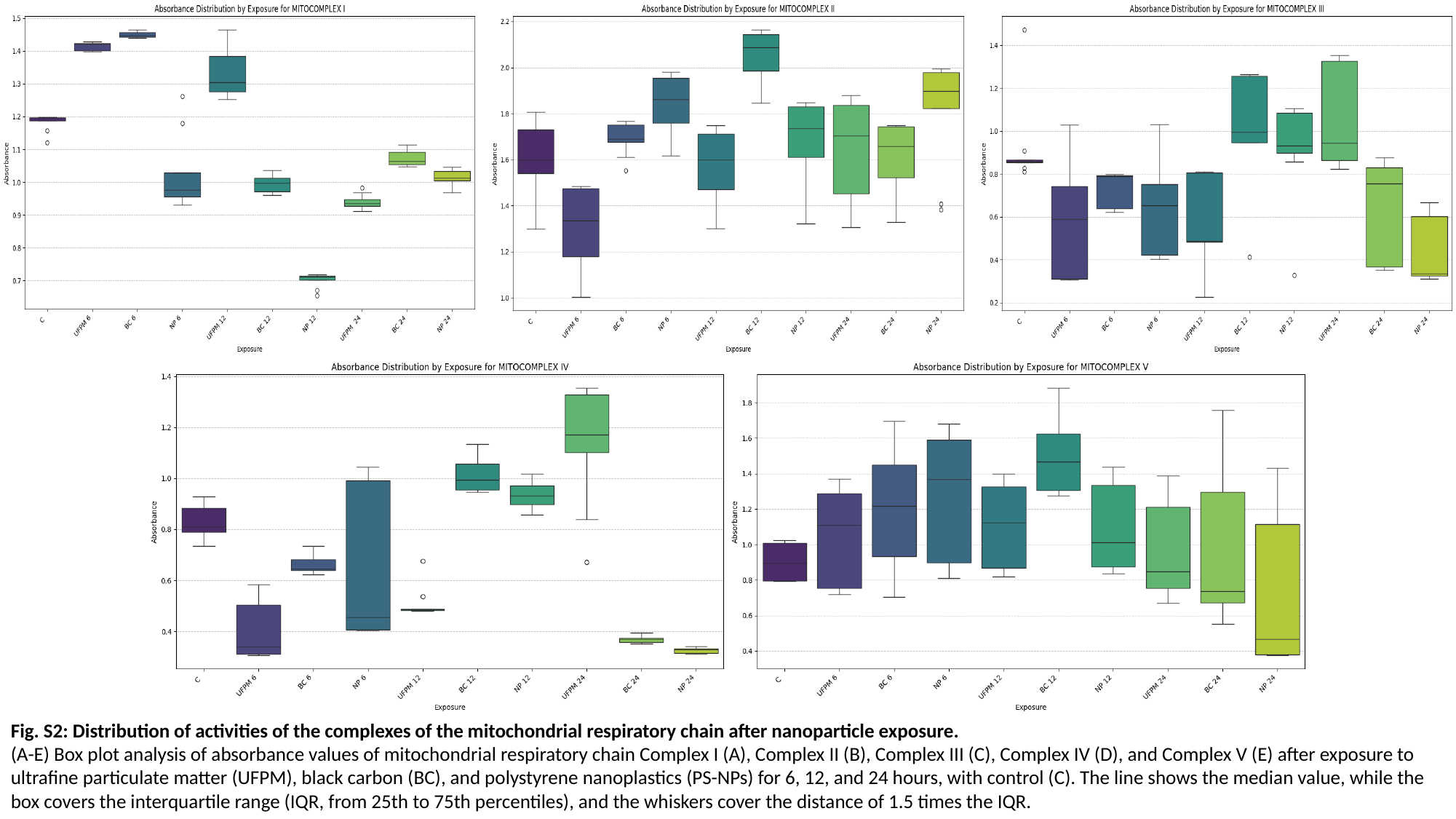

Fig. S2: Distribution of activities of the complexes of the mitochondrial respiratory chain after nanoparticle exposure.
(A-E) Box plot analysis of absorbance values of mitochondrial respiratory chain Complex I (A), Complex II (B), Complex III (C), Complex IV (D), and Complex V (E) after exposure to ultrafine particulate matter (UFPM), black carbon (BC), and polystyrene nanoplastics (PS-NPs) for 6, 12, and 24 hours, with control (C). The line shows the median value, while the box covers the interquartile range (IQR, from 25th to 75th percentiles), and the whiskers cover the distance of 1.5 times the IQR.
